# SLC13A2-transported citrate is a metabolic signal for PKM2 acetylation and degradation to suppress tumor growth

**DOI:** 10.1101/2024.05.04.591017

**Authors:** Mengyao Qin, Li Shi, Hao Chen, Chan Liu, Zhiquan Tang, Donghao An, Wanting Yu, Dandan He, Chang Shao, Shengtao Yuan, Hong Yu, Haiping Hao, Jing Xiong

## Abstract

**Background and aims:** Metabolic reprogramming represents as a pivotal hallmark for cancer, but TCA cycle in tumorigenesis and progression has long been neglected. Solute carrier (SLC) transporters mediate the transport of TCA cycle intermediates across membrane, but their functions in cancer pathogenesis remains unclear.

**Approaches & Results:** Using integrated analysis of solute carrier (SLC) transporters for TCA cycle intermediates, we found that SLC13A2 was consistently downregulated in hepatocellular carcinoma (HCC) cells and liver tissues from human patients and heterogeneous mouse models. Adeno-associated virus (AAV)-transduced liver-specific knockout or overexpression of SLC13A2 promoted or ameliorated HCC progression in the primary mouse model, demonstrating that SLC13A2 served as a protective factor during HCC pathogenesis. SLC13A2 inhibited HCC cell proliferation by decreasing mitochondrial function via suppressed oxygen consumption and ATP production. Combined with metabolic flux analysis, we found that SLC13A2 imported citrate, which secreted acetyl-CoA as a precursor for the acetylation of pyruvate kinase muscle isozyme M2 (PKM2), which led to its protein degradation. Decreased activity of pyruvate kinase depleted pyruvate for the TCA cycle, thus inhibiting amino acid synthesis and nucleotide metabolism. Additionally, a decrease in nuclear PKM2 protein transduced to reprogrammed gene transcription for cell proliferation and metabolism which is required for tumor growth.

**Conclusions:** This study revealed that citrate transported by SLC13A2 acts as a signal to disrupt metabolic homeostasis for tumor growth and suggests potential drug targets for HCC therapy.

**graphic abstract:** 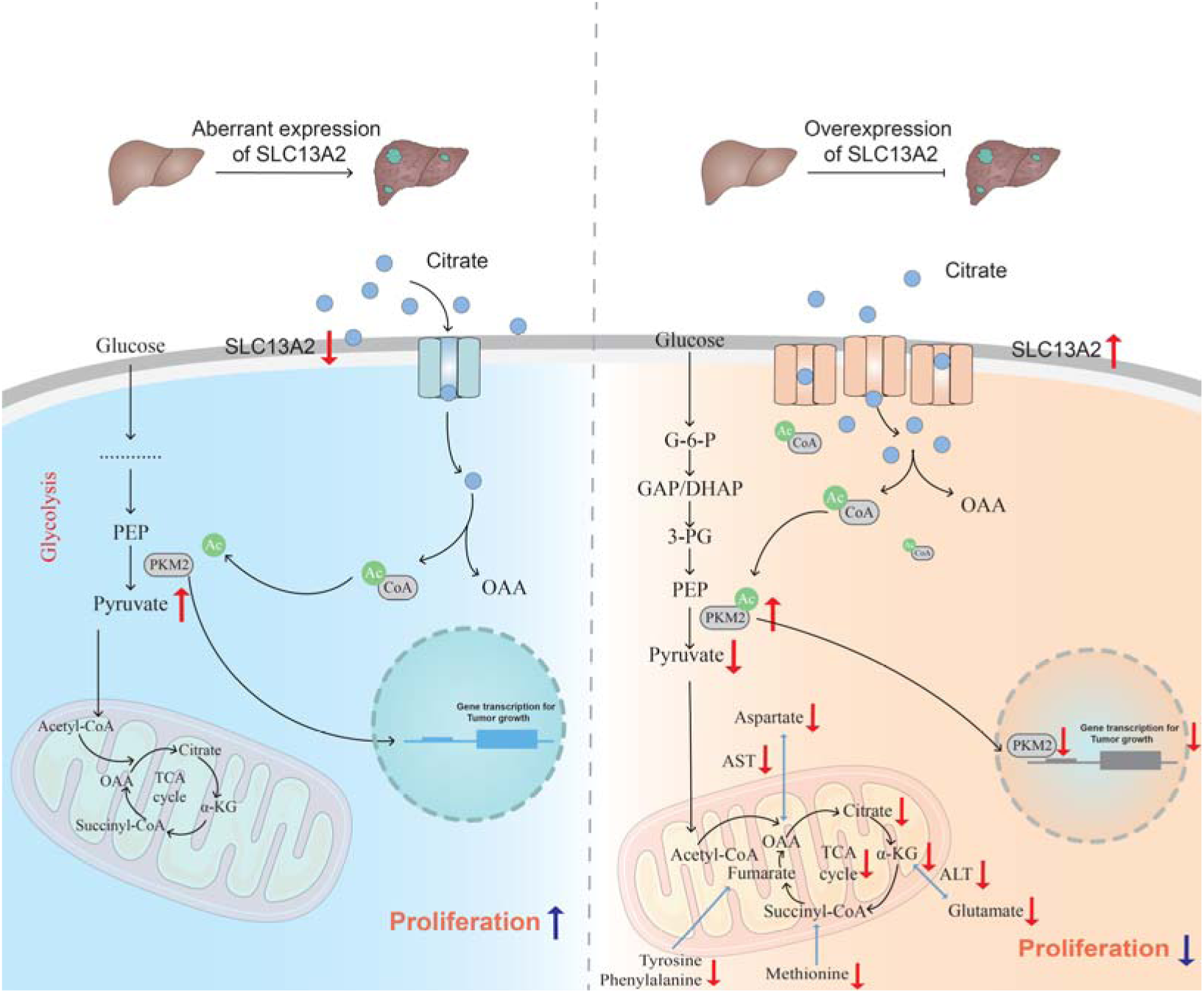

## Introduction

Cancer is characterized by metabolic reprogramming to fulfill its increased demands for uncontrolled cell proliferation. Malignant cells, including hepatocellular carcinoma (HCC), rapidly take up and breakdown nutrients, which contributes to cancer progression (1). Accumulating evidence indicates that elucidating the metabolic dysregulation of tumor cells provides an exciting new direction for targeted HCC therapy (2).

The tricarboxylic acid (TCA) cycle serves as a convergence point for nutrient metabolism and macromolecule synthesis (3). It completely catabolizes many types of fuel sources from the diet, including glucose, amino acids and fatty acids, and then provides precursors for building blocks in macromolecule synthesis, such as nonessential amino acids, purines and pyrimidines (4). In normal cells, glucose produces pyruvate, which enters the TCA cycle and acts as a major source of adenosine triphosphate (ATP) through oxidative phosphorylation. In contrast, cancer cells often truncate glucose metabolism away from the TCA cycle through aerobic glycolysis and therefore are more addicted to glutamine or rely on fatty acids to replenish TCA cycle intermediates (5). Aberrant TCA function due to somatic/germline enzyme mutations or altered activity, such as mutation of isocitrate dehydrogenase 2 (mIDH2) and increased activity of citrate synthase (CS), has been implicated in cancer pathogenesis (4). Moreover, changes in the expression of intermediate transporters have been shown to rewire the metabolism of cancer cells (6). Both oncogenes and tumor suppressors manipulate TCA cycle flux by regulating the expression of intermediate transporters and the activity of enzymes (4). In light of the well-accepted concept that cancer cells mainly obtain energy through aerobic glycolysis, the role of the TCA cycle in cancer metabolism and tumorigenesis has long been neglected and deserves further elucidation.

To ensure the function of the TCA cycle, the concentrations of intermediates are precisely regulated by the expression of transporters across the plasma membrane and mitochondrial inner membrane (7). Solute carrier (SLC) transporters, which are highly expressed in the liver, represent the second largest group of membrane proteins after G protein-coupled receptors (8). SLCs mediate the transmembrane transport of various types of nutrients and metabolites, such as glucose, amino acids, lipids and nucleotides, and thus serve as metabolic switches in cells (9). As most SLCs transport small molecules, they are considered promising targets for drug development (10). Some SLC transporters, such as SLC1A5 inhibitor, have been successfully targeted for the treatment of cancer (11, 12). There are six main SLC families with members identified as carriers that regulate TCA cycle flux by transporting TCA cycle intermediates such as citrate, succinate and malate through the plasma membrane or mitochondrial matrix according to an updated version of the guide to transporters (13). The role of these SLC transporters in the regulation of TCA cycle function and tumor growth is largely elusive. Based on previous findings, it is speculated that these SLC family members control anaplerotic or cateplerotic flux in response to metabolic adaptation of cancer cells.

Here, we identified SLC13A2 (also known as Na^+^-dicarboxylate cotransporter 1, NaDC1) as a downregulated SLC transporter for TCA cycle intermediates in HCC liver tissues and cells. By examining a liver-specific SLC13A2 knockout/overexpression mouse HCC model, we further identified SLC13A2 as a tumor suppressor for HCC progression. Combined with metabolomics and transcriptomics analysis, SLC13A2, a carrier for citrate, depletes TCA cycle intermediates, suppresses the biosynthesis of macromolecules, and remodels gene transcription for tumor growth. Our findings imply that citrate acts as a metabolic signal transported by SLC13A2, which has the potential to provide an attractive strategy for treating HCC.

## Materials and methods

### Animal studies

All animal experiments were approved by the IACUC (Institutional Animal Care and Use Committee) of China Pharmaceutical University (Approval number: 2020-08-006). Male C57BL/6J mice aged 6-8 weeks were purchased from the University of Yangzhou and randomly grouped. Mice were injected with oncogene plasmids through the tail vein for 4-7 s via a technique called hydrodynamic transfection with oncogene delivery and Sleeping Beauty (SB)-mediated somatic integration for stable and long-term gene expression in hepatocytes, as previously described (14). The oncogenic plasmids used were pT3-EF1aH-c-MET (20 μg) and pT3-EF1aH-ΔN90-β-catenin (20 μg), along with SB transposase (5 μg). The oncogene plasmids were diluted with saline to 10% body weight (2 ml/20 g). For liver-specific SLC13A2 knockout experiments, single-guide RNA (sgRNA) sequences were designed based on a CRISPR design tool targeting the first coding exon (Table S1) and constructed together with thyroid hormone-binding globulin (TBG)-Cre into adeno-associated virus 8 (AAV8, 5 × 10^11^ genome copies/mouse), which was injected into the tail vein of *Rosa26-flox-STOP-flox-Cas9* knockin mice (*Rosa26-LSL-Cas9*, Jackson, #024857) as previously described (15). For liver-specific SLC13A2 overexpression experiments, adult male mice (6-8 weeks of age, ∼20 g body weight) were injected with AAV8 (1 × 10^11^ genome copies/mouse) under the control of the liver-specific TBG promoter via the tail vein (AAV-TBG-GFP versus AAV-TBG-SLC13A2). The plasmids were constructed using the primers shown in Table S1 and used for the AAV8 package. All sufferings of the animal are minimized during the experiments.

### Human studies

Human HCC liver samples were obtained during partial liver resection surgery at Jiangsu Taizhou People’s Hospital (Taizhou, Jiangsu, China) under a protocol approved by the Medical Ethics Committee of Taizhou People’s Hospital (Approval number: 2022-03-017). The study was conducted in accordance with the ethical requirements of the Declaration of Helsinki. Fresh liver tissues from 8 HCC patients (including cancerous tissues and adjacent paracancerous tissues) were obtained during surgical resection at the Hospital of Jiangsu Taizhou People (Taizhou, Jiangsu, China). Both male and female individuals were included. The average age of the patients was 68.125 years. Written informed consent was provided by all participants.

### Cell culture

All cell lines were characterized by short tandem repeat profiling. The human HCC cell line HepG2 was obtained from the Institute of Biochemistry and Cell Biology of the Chinese Academy of Sciences (Shanghai, China). The mouse HCC cell line Hepa1-6 was obtained from the American Type Culture Collection (CRL-1830) as described previously (16). Mouse primary hepatocytes were isolated from wild-type male C57BL/6J mice and cultured as described previously (16).

### Quantitative reverse transcriptase□polymerase chain reaction (qRT□PCR)

Total RNA was extracted from cells or liver tissues using TRIzol reagent (Vazyme, Nanjing, China). cDNA was synthesized using the PrimeScript™ RT Reagent Kit (Vazyme, Nanjing, China). qRTLPCR was performed with the qPCR SYBR Green Master Mix (High ROX Premixed) kit (AG Bio, Hunan, China). The primers designed with NCBI primer BLAST are shown in Table S1.

### Immunoblotting and immunoprecipitation

Protein samples were mixed with 3 × SDS loading buffer, separated using SDSLPAGE, and then transferred onto a polyvinylidene fluoride membrane. The membranes were incubated with primary antibodies at 4°C for 12-16 h and then with secondary antibodies for 1 h at room temperature. Enhanced chemiluminescence reagents were used to visualize the target proteins. For the immunoprecipitation assay, cell extracts were incubated with primary antibodies or control IgG in a rotating incubator overnight at 4°C, followed by incubation with protein A agarose beads (Beyotime, Shanghai, China) for another 4 h. The immunoprecipitates were washed five times with lysis buffer and analyzed by immunoblotting. The primary and secondary antibodies used are shown in Table S2.

### Histological analysis and immunohistological chemistry staining

Formalin-fixed and paraffin-embedded mouse liver sections were stained with hematoxylin and eosin (H&E). For immunohistological chemistry staining, the samples were dewaxed and hydrated. The antigen was retrieved with citric acid buffer (pH 6.0), blocked with horse serum, and incubated with Ki67 (Abcam, ab279653) antibody at 4°C for 12-16 h and then with a secondary antibody conjugated with horseradish peroxidase for 1 h at room temperature. Images were taken with a BX53 light microscope (Olympus, Japan).

### Plasmid construction

The plasmids PLVX-GFP-N1 and PLVX-GFP-SLC13A2 were constructed using the primers listed in Table S1. An empty control vector for pReceiver-M02 (EX-NEG-M02) and a mouse PKM2 expression plasmid (EX-Mm04490-M02) were purchased from Guangzhou FulenGen Co., Ltd. (Guangzhou, China).

### siRNA and plasmid transfection

For siRNA or plasmid transfection, Hepa1-6 cells were seeded at 40% or 80% confluence the next day and transfected with siRNAs or plasmids using Lipo8000 (Beyotime, Shanghai, China) according to the manufacturer’s instructions (siRNA sequences are shown in Table S1). HepG2 cells were infected with adenovirus (pAdeno-MCMV-3XFlag versus pAdeno-MCMV-SLC13A2-3XFlag, OBiO, Shanghai, China) at an MOI of 25. After 12-18 h of incubation, the medium was changed to complete medium, and the cells were then cultured for the indicated period before the experiments.

### Assays for cell proliferation *in vitro*

For the knockdown experiment, Hepa1-6 cells were transfected with siRNAs for 24 h, digested for seeding in 6-well plates at a density of 2.5 × 10^3^ cells/well, and then cultured for 7 days. For the overexpression experiment, Hepa1-6 and HepG2 cells were transfected with the overexpression plasmid or adenovirus, respectively, for 24 h, digested for seeding onto a 6-well plate at a density of 1 × 10^3^ cells/well, and cultured for 14 days. Cell colonies were fixed with 4% formalin for 15 min and stained with 0.4% crystal violet for 10 min. For the 5-ethynyl-2’-deoxyuridine (EdU) incorporation assay, Hepa1-6 and HepG2 cells were incubated with EdU for 4 h, fixed for 30 min and analyzed using a commercial kit (Beyotime, Shanghai, China). Similarly, cell cycle arrest and cell viability were analyzed using a cell cycle analysis kit (Beyotime, Shanghai, China) and a CCK8 assay kit (Vazyme, Nanjing, China), respectively, according to the manufacturer’s instructions.

### Measurement of the oxygen consumption rate (OCR)

The cells were seeded into the wells of a Seahorse XF 96-well culture plate (Agilent, Japan) at a density of 8 × 10^4^ cells/well and cultured overnight at 37°C and 5% CO2. Six wells of cells from each group were used for the analysis. The OCR was measured with the assurance that the cells were even and approximately 90% confluent in a monolayer. The sensor cartridge was incubated in a non-CO_2_ incubator at 37°C overnight. Cells in growth media were replaced with fresh and prewarmed XF media (10 mM glucose and 1 mM sodium pyruvate, 1 mM glutamine) and then incubated in a non-CO_2_ incubator at 37°C for 45 min. Compounds were diluted with fresh XF media and injected for measurement as follows (oligomycin 1.5 μM, FCCP 1 μM and rotenone/antimycin 0.5 μM). The data were analyzed directly by Seahorse software. The basal and maximal OCRs were obtained for statistical analysis.

### Mitochondrial membrane potential assay

Hepa1-6 cells were seeded at a density of 5 × 10^4^ cells/well in a 24-well plate. After transfection with the plasmid for 48 h, the cells were stained with JC-1 according to the manufacturer’s instructions (Beyotime, Shanghai, China). Images were taken using an inverted fluorescence microscope (IX71, Olympus, Japan) with the imaging software Olympus cellSens Standard software.

### Isolation of subcellular fractions

The procedures were conducted on ice with prechilled buffers and equipment. Hepa1-6 cells in 10 cm dishes were transfected with adenovirus and harvested after 72 h. After being washed with PBS three times, the cell pellets were suspended in homogenization buffer (225 mM D-mannitol, 75 mM sucrose, 0.1 mM EDTA, 30 mM Tris-HCl, pH 7.5) and then transferred to a 2-mL Dounce homogenizer to disrupt the cells with several strokes of the pestle. For each stroke, the pestle was pressed straight down the tube to maintain a firm and steady pressure. When checking the degree of homogenization with a phase-contrast microscope, eight to nine naked nuclei for every whole cell indicated a good result. The homogenate was transferred to a centrifuge tube and spun down at 500 × g for 5 min. While the supernatant was used for preparing the cytoplasm at 7000 × g for 10 min, the pellet was gently suspended in homogenization buffer followed by centrifugation at 700 × g for 10 min to collect the nuclei. The extracted subcellular fractions were confirmed by immunoblotting for cytosolic and nuclear P-tubulin and PCNA, respectively.

### ATP measurement

Transfected cells were seeded into 96-well culture plates at a density of 2 × 10^3^ cells/well. After 72 h, ATP production was detected with a CellCounting-Lite^®^ 2.0 Luminescent Cell Viability Assay kit (Vazyme Biotech, Nanjing, China) by a plate reader (Beckman Coulter, Krefeld, Germany) according to the manufacturer’s instructions.

### Metabolomic analysis

Hepa1-6 cells were collected, and metabolites were extracted after adenovirus transfection for 72 h. Cell metabolites were dissolved in 0.5 mL of ice-cold 80% methanol solution containing 1 μg/mL 4-chloro-DL-phenylalanine (Sigma, C6506). The samples were then incubated for 30 min in a −80°C freezer, followed by centrifugation at 13,000 ×Lg for 15 min at 4°C, after which the supernatant was dried using a vacuum centrifugal concentrator (Thermo, Massachusetts, USA). Before analysis, the samples were reconstituted with methanol, followed by centrifugation at 18,000 ×Lg for 10 min at 4°C twice, and untargeted metabolomics was conducted using a 6546 LC/Q-TOF (Agilent, USA) instrument equipped with an electrospray ionization (ESI) source. Separation was achieved on an XBridge Premier BEH Amide VanGuard FIT column (2.5 μm, 4.6 mm × 150 mm, Waters, USA). MS data were acquired in negative ESI mode over a range of m/z 50 to 850. The mobile phase consisting of water (containing 0.3% ammonium acetate and 0.3% ammonia in phase A, v/v) and acetonitrile (phase B) was delivered at a flow rate of 0.4LmlLmin^−1^. The total elution time was 36 min for the gradient program. The data were analyzed by Progenesis QI with conditions of P<0.05, fold change>2, and VIP>1 to screen for differentially abundant metabolites. Metabolic pathway analysis was performed based on the MetaboAnalyst 5.0 platform.

For the quantitative determination of metabolites, cell samples and mixed standards of gradient concentrations were measured on a TripleTOF 5600 system (Sciex, USA) by direct infusion. The instrument was set to acquire data over a m/z range of 100-2,000 for the TOF-MS/MS scan. Aliquots of reconstituted supernatants (5 μL) were analyzed with an XBridge BEH Amide column (3.5 µm, 4.6LmmL×L100Lmm, Waters, USA) on an LC-30 HPLC system (Shimadzu). The mobile phase consisting of water/acetonitrile (95:5, v/v) (containing 0.1% ammonium acetate and 0.3% ammonia in phase A, v/v) and acetonitrile (phase B) was delivered at a flow rate of 0.4LmlLmin^−1^. The total elution time was 36 min for the gradient program. The concentrations of the tested substances were calculated by comparing the parent ion and daughter ion as well as the retention time with the standard curve formed by the gradient standards.

### Stable isotope resolved metabolomics (SIRM) analysis

The culture medium was replaced by U-13C^6^-glucose (Cambridge) in glucose-free DMEM (Sigma) after the transfection of Hepa1-6 cells for 60 h. The cells were labeled at the indicated time points. An unlabeled culture was prepared in parallel by adding equal concentrations of unlabeled glucose to the media to identify unlabeled metabolites. Cell metabolites were extracted with 80% methanol and completely concentrated to dryness using a Speedvac (Thermo Fisher Scientific, USA). The samples were redissolved in pure methanol and used for LCLMS/MS analysis. Standard metabolites were observed, and the MS/MS spectra of these compounds were manually confirmed according to Metlin (www.metlin.scripps.edu) or HMDB (www.HMDB.ca).

### RNA-seq and bioinformatic analysis

Liver RNA sequencing was performed on DNBSEQ-T7 cells from GFP- or SLC13A2-OE mice in the HTVi model. The fastq files were aligned to the mouse reference genome (mm10) using the aligner STAR. RNA-seq fastq data have been deposited into the Sequence Read Archive (PRJNA1083485). The published RNA-seq raw data for heterogeneous liver cancer induced by diverse cancer driver genes were downloaded from the NCBI Sequence Read Archive with accession code PRJNA674008 (17). The published RNA-Seq raw data of human HCC cancerous and paracancerous specimens were downloaded from the Gene Expression Omnibus database (GSE193567) (18). TCGA HCC RNA-seq data were downloaded from the NIH-GDC repository (https://portal.gdc.cancer.gov/), which included 371 primary HCC patients in level 3. The HCC samples were aligned according to the expression level of PKM2, where top quarter considered as PKM2-high group was compared with bottom quarter as PKM2-low group to screen out PKM2-regulated gene cluster in clinical datasets. Differential gene expression analysis was performed using the Deseq2 package in the R program. The ENSEMBL numbers in the analysis results were converted to gene names using probes in the human gene annotation package, and genes with a log2FoldChange greater than 1 or −1 and a P value less than 0.05 were selected for plotting using the ggplot2 package. Heatmaps and volcano plots were generated by R. Gene enrichment analysis was performed using MetaScape (www.metascape.org). The overall differentially expressed genes and the upregulated and downregulated genes in SLC13A2-OE mice were independently subjected to motif enrichment analysis (HOMER, findMotif.pl).

### Enzymatic activity assay

Approximately 100 mg of liver tissue was ground with a 1:9 weight/volume ratio of PBS and centrifuged at 2500 × g at 4°C for 10 min. Hepa1-6 cells were transfected in 6-well plates with plasmids for 72 h. The collected cells were mixed with 500 μL of PBS and homogenized manually. The enzymatic activity of the homogenate was detected by alanine transaminase (ALT) and aspartate transaminase (AST) kits (Jiancheng, Nanjing, China). Protein concentrations were quantified using the Bradford protein assay and used for the normalization of enzymatic activity.

### Cycloheximide (CHX) chase assay

To assess the half-life of the proteins, Hepa1-6 cells overexpressing SLC13A2 were treated with 30 μg/mL CHX, a protein synthesis inhibitor, and harvested at 0 h, 3 h, 6 h or 9 h for immunoblot analysis. The densitometry of the bands was quantified using ImageJ.

### Statistical analysis

The data are expressed as the mean ± SEM. The differences between two groups were statistically analyzed using a two-tailed Student’s *t* test. Differences from the growth curve study were analyzed by two-way ANOVA, followed by Bonferroni multiple comparisons. P<0.05 was considered to indicate statistical significance.

## Results

### SLC13A2 is a downregulated SLC transporter for TCA cycle intermediates in HCC

To explore the formerly overlooked role of the TCA cycle, we focused on transporters of TCA cycle intermediates in HCC. To date, SLC families, including SLC5, SLC13, SLC16, SLC25, SLC33 and SLC54, have been identified to transport TCA intermediates such as citrate, succinate, a-KG, malate, L-lactate and pyruvate, according to an updated version of the guide to transporters (13) (Figure 1A). We then analyzed RNA-seq data from heterogeneous liver tumors induced by diverse cancer driver genes selected by mutational frequency from human HCC cohorts with different etiological factors and ethnic origins (17). SLC13A2 was potently and consistently downregulated in all liver tumors with obviously heterogeneous phenotypes, showing an extraordinary pattern compared with that of other detectable SLC transporters (Figure 1B).

**Figure 1.**
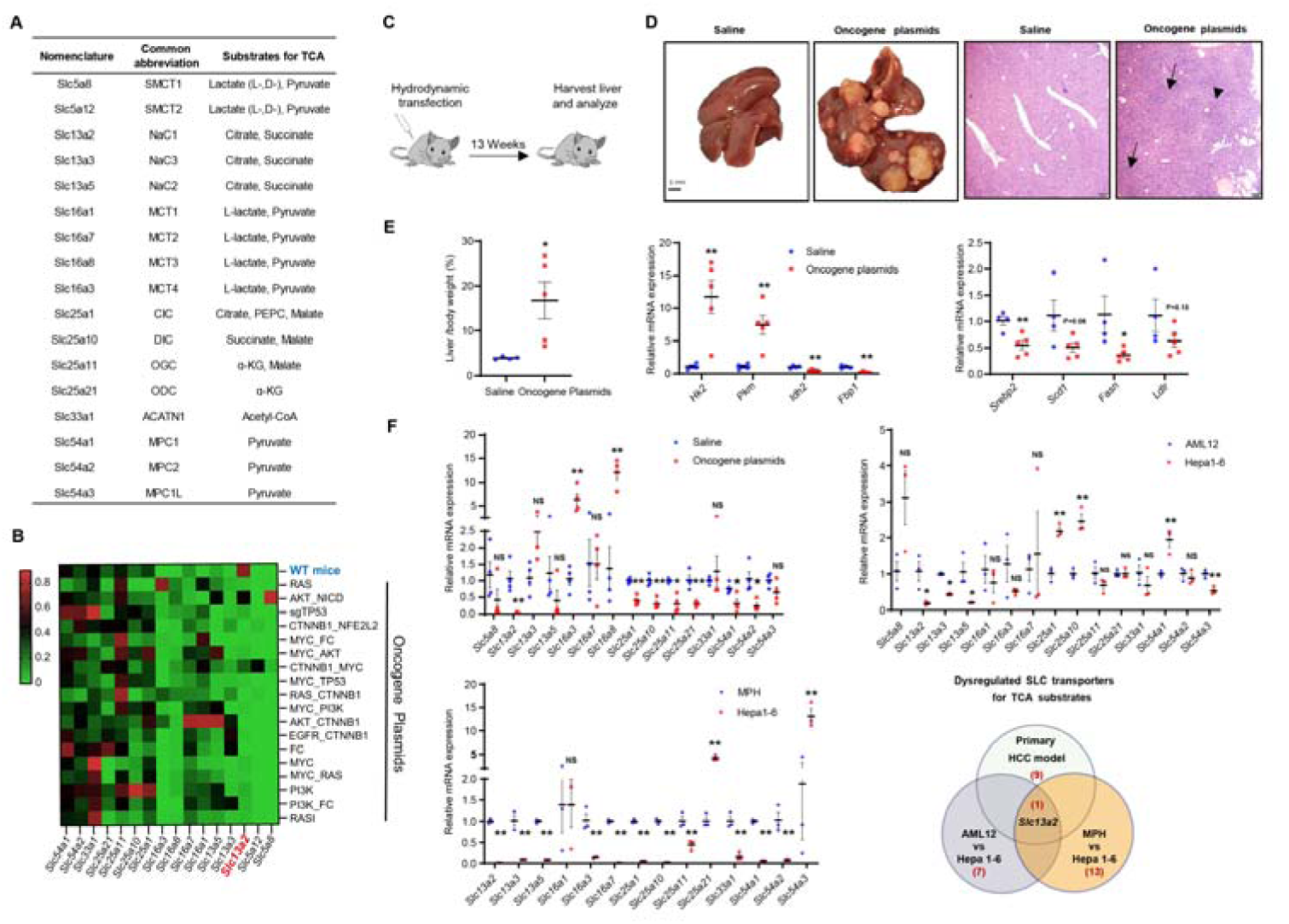
SLC13A2 is a transporter for TCA cycle intermediates deregulated in HCC. **A.** The SLC transporters for TCA cycle intermediates are summarized. **B.** Heatmap of detectable SLC transporters for TCA cycle intermediates in heterogeneous primary liver cancer (PLC) models established using hydrodynamic tail vein injection (HTVi)-generated genome editing of cancer driver genes selected by mutational frequency from human HCC cohorts (SRA accession code PRJNA674008). **C.** Schematic showing the study design of the HTVi model by injecting sleeping beauty (SB) transposase plasmids combined with c-Met and ΔN90-β-catenin oncogene plasmids (c-MET: ΔN90-β-catenin:SB = 20 μg:20 μg:5 μg). **D.** Representative liver morphology after dissection (left, Scale bar: 5 mm) and histological staining (right, Scale bar: 200 µm). The arrows indicate areas with densely packed and multinucleated hepatocytes with an increased nucleo-plasma ratio. **E.** Liver/body weight ratio (left) and mRNA expression of metabolic genes in liver tissues of the HTVi model (right) (n=4 for the control group and n=5 for the HCC model group). **F.** Methodology to screen candidate SLC transporters for HCC progression based on the relative expression of SLC transporters in the HTVi model, mouse HCC cells, AML12 cells and mouse primary hepatocytes (MPHs). The SLC transporters that are detectable for TCA intermediates in liver tissues of the HTVi model (n=4 for the control group and n=5 for the HCC model group), AML12 cells, primary hepatocytes and mouse HCC Hepa1-6 cells were validated by qPCR (n=3 independent experiments). The data are presented as the means ± SEMs. *P < 0.05, **P < 0.01, ns = not significant; two-tailed unpaired Student’s *t* test.

Based on the above data, we generated a primary HCC mouse model (HTVi model) to evaluate the changes in the above SLC family members via hydrodynamic transfection of oncogene plasmids in combination with SB transposase (Figure 1C). Compared with those in the saline group, obvious tumor nodules were observed in the livers of the mice that were transfected with the oncogene plasmids, as shown by the presence of densely packed and multinucleated hepatocytes, as well as an increased nucleoplasmic ratio according to histological staining (Figure 1D). The liver/body weight ratio significantly increased in the oncogene-transduced group (Figure 1E, left). The mRNA expression of genes involved in glucose metabolism and fatty acid metabolism enzymes in the oncogene plasmid group was consistent with the metabolic adaptation of tumor cells (Figure 1E, right). These findings suggested that the primary HCC mouse model was generated successfully.

Next, we detected the expression of SLC transporters of interest in liver tissues and found that the expression of *Slc16a3* and *Slc16a8* significantly increased, while that of *Slc13a2, Slc25a1, Slc25a10, Slc25a11, Slc25a21, Slc54a1,* and *Slc54a2* significantly decreased in liver tissues from mice with HCC (Figure 1F top left). Among these genes, *Slc13a2* decreased 10-100 times and attracted our attention because it exhibited the most abundant variation. Moreover, the expression of SLC transporters was evaluated in mouse HCC Hepa1-6 cells and mouse primary hepatocytes, and the results showed that the expression of *Slc25a21* and *Slc54a3* significantly increased, while that of other SLC transporters of interest decreased, except for *Slc16a1* (Figure 1E bottom left). Similarly, we detected SLC transporter expression in mouse HCC Hepa1-6 cells compared with that in immortalized mouse hepatocyte AML12 cells. The gene expression of *Slc25a1*, *Slc25a10 and Slc54a1* increased, while that of *Slc13a2*, *Slc13a3*, *Slc13a5*, and *Slc54a3* decreased in the HCC cell line (Figure 1E, upper right). Among the above SLC transporters, *Slc5a12* was undetectable in both HCC liver tissues and cell lines. Using integrated analysis of SLC expression in primary HCC liver tissues and cell lines, we found that *Slc13a2* was the only gene consistently deregulated both *in vivo* and *in vitro* (Figure 1F bottom right). In addition, reduced protein expression of SLC13A2 was further confirmed in HCC liver tissues, while the levels of proteins and cell signaling pathways that promote cell proliferation were significantly increased (Figure 2A). These results indicate that SLC13A2 is downregulated in HCC and stands out from the other SLC transporters for TCA cycle intermediates.

**Figure 2.**
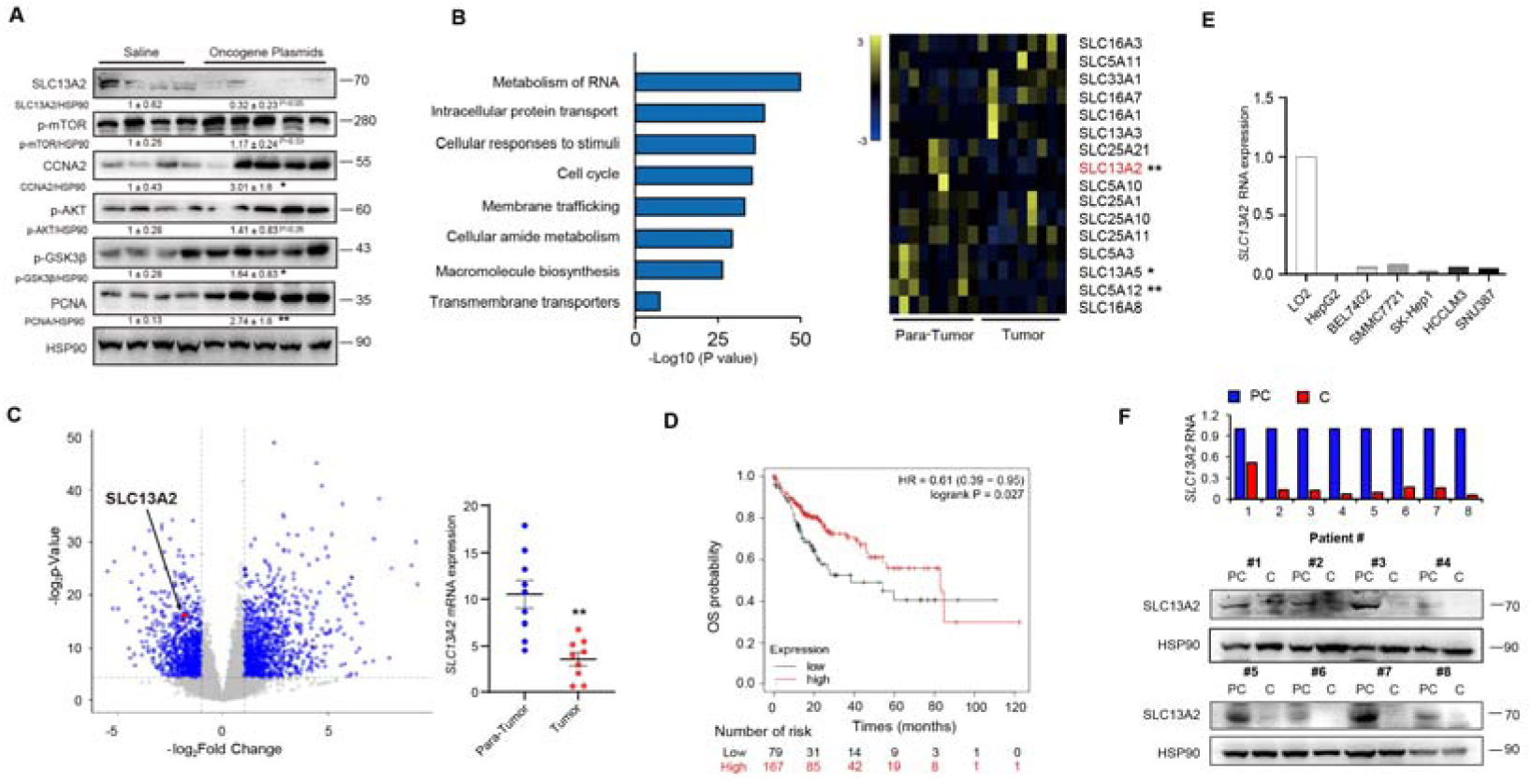
SLC13A2 expression is decreased in human HCC specimens and cell lines, which is consistent with the findings in animal models. **A.** Immunoblots of total liver lysates from the HTVi model. **B-C.** Analysis of published RNA-Seq data of human HCC cancerous and paracancerous specimens from GEO datasets (GSE193567). Ontology enrichment analysis of differentially expressed genes (B, left). Heatmap showing the detectable SLC transporters for TCA cycle intermediates (B, right). Volcano plots showing DEGs with emphasis on SLC13A2 (C, left). The expression level of SLC13A2 in tumor tissues compared with that in para-tumor tissues (C right). The data are presented as the mean ± SEM (n=9). **P < 0.01, tumor versus para-tumor tissues, two-tailed unpaired Student’s *t* test. **D.** Kaplan–Meier (KM) analysis of survival in male patients with HCC stratified by high or low SLC13A2 expression is shown. **E.** The RNA expression of SLC13A2 in human HCC cell lines versus L02 cells after being split and cultured for 48 h. **F.** The RNA and protein expression of SLC13A2 was analyzed in eight clinical HCC patients. PC, paracancerous; C, cancerous.

By analyzing a published Gene Expression Omnibus dataset (GES193567), we identified 4247 differentially expressed genes (DEGs) between cancerous and paracancerous tissues of human HCC patients. Gene enrichment analysis revealed that the DEGs were enriched in macromolecular biosynthesis, membrane trafficking and transmembrane transporters (Figure 2B, left). In this dataset, SLC13A2 was identified as one of the three significantly altered SLC transporters for TCA cycle intermediates between paratumor and tumor tissues (Figure 2B, right). As shown for all DEGs, SLC13A2 was significantly reduced in tumor tissues (Figure 2C). Moreover, Kaplan–Meier survival curves revealed that male patients with high SLC13A2 expression had a better prognosis and a greater overall survival rate (Figure 2D). Additionally, SLC13A2 mRNA was significantly lower in human HCC cell lines than in the normal human liver cell line LO2 (Figure 2E). Furthermore, we collected liver tissues from clinical patients with HCC and analyzed the expression of SLC13A2. We confirmed that all HCC patients had lower SLC13A2 expression in cancerous tissues than in paracancerous tissues (Figure 2F). The above data suggest that reduced expression of SLC13A2 is conserved in mouse and human HCC and is positively associated with the prognosis of this disease.

### SLC13A2 inhibits HCC progression in a mouse model induced by hydrodynamic transfection of oncogenes

The function of SLC13A2, a previously described transporter in renal tubules and intestinal cells, in physiological homeostasis in the liver is still obscure. Initially, we studied the role of SLC13A2 in the liver homeostasis without any insults via hepatocyte-specific SLC13A2 knockout or overexpression by AAV-mediated transduction as described previously (**EMBOJ-2024-118432)**. The unchanged blood glucose, body weight, liver weight, histological staining and the expression of metabolic genes in the settings of SLC13A2 manipulation imply that hepatocyte-specific knockout or overexpression of SLC13A2 primes the cell metabolic status for HCC tumorigenesis but is not sufficient to directly induce the initiation of HCC.

To explore the effect of SLC13A2 on the tumorigenesis and progression of HCC, we first used CRISPR/Cas9-mediated LKO mice. The mice were injected with oncogene plasmids using the HTVi model after 2 weeks of AAV8 injection and sacrificed after 8 more weeks (Figure 3A). As expected, the knockout efficiency of SLC13A2 was verified at the mRNA and protein levels (Figure 3B). Although there were no significant differences in body weight or blood glucose between the two groups, the liver weight/body weight ratio significantly increased in the LKO group (Figure 3D). The number of surface tumor nodules was significantly greater in LKO mice than in control mice (Figure 3C). While the percentage of tumor incidence was similar between the two groups, their tumor sizes were significantly different. The SLC13A2 LKO group had remarkably more mice with larger maximal tumor sizes than did the control group (Figure 3E). As indicators of liver tumor burden (19), the enzymatic activity of serum and liver ALT/AST also increased with SLC13A2 ablation (Figure 3F). Histological staining revealed larger nodules with more irregularly shaped nuclei in the liver H&E-stained slides from the SLC13A2 LKO group (Figure 3G). Immunohistochemical staining of Ki67 also revealed 16.6% nuclear positivity in the control group and approximately 22.9% nuclear positivity in the SLC13A2 LKO group, demonstrating a significant increase in cell proliferation induced by the knockout of SLC13A2 *in vivo* (Figure 3H). Consistently, the phosphorylation of mTOR, AKT and GSK3β, which are involved in cellular signaling related to cell proliferation and metabolism, was significantly elevated, consistent with the promotion of tumor growth in liver-specific knockout mice (Figure 3I).

**Figure 3.**
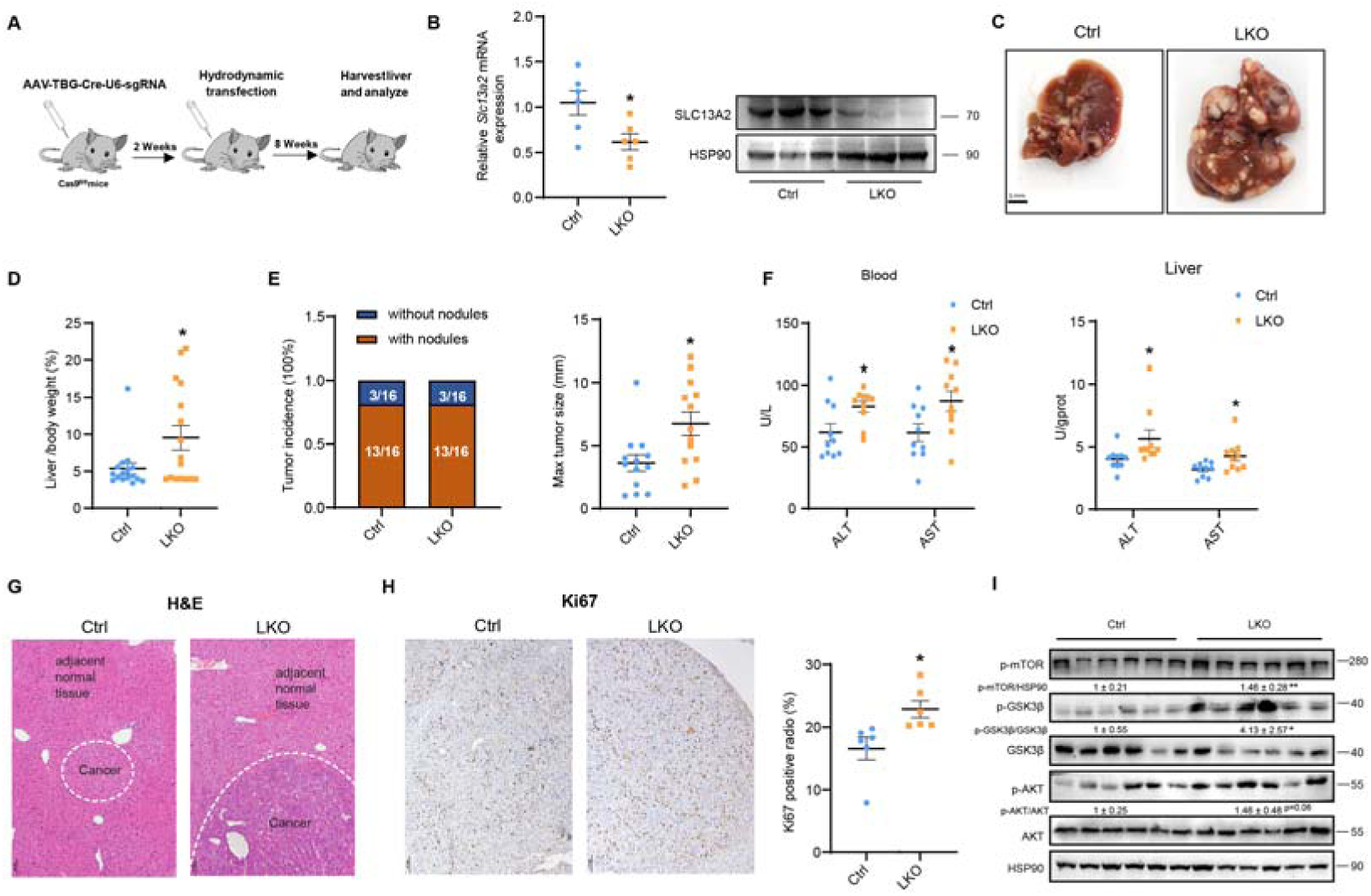
Liver-specific knockout of SLC13A2 promotes HCC tumor growth. **A.** A diagram of Cas9-mediated liver-specific knockout (LKO) of SLC13A2 and study design. AAV-TBG-Cre-U6-Lacz (Ctrl) and AAV-TBG-Cre-U6-sgRNA (LKO) viruses (5 × 10^11^ VP/mouse) were injected into *Rosa26-FSF-Cas9* knock-in mice via the tail vein. The transduced mice were subjected to the HTVi model by injecting SB in combination with oncogene plasmids (c-MET: ΔN90-β-catenin: SB = 20 μg:20 μg:5 μg). **B.** qPCR analysis (left, n=6 per group) and immunoblots (right) showing the SLC13A2 knockout efficiency in mouse liver tissues. **C.** Representative liver morphology after dissection. Scale bar: 5 mm. **D.** Liver/body weight ratio. n=16 per group. **E.** Tumor incidence showing the number of mice with or without nodules (left) and the maximal tumor size (right, n=13 per group) in the two groups. **F.** Liver ALT/AST and serum ALT/AST activity. **G.** Representative images of H&E staining. **H.** Immunohistochemical staining for Ki67. The Ki67-positive ratio was calculated (n=6 per group). **I.** Immunoblots of total liver lysates showing cell signaling for tumor growth. The data are presented as the means ± SEMs. *P < 0.05, LKO vs Ctrl group, two-tailed unpaired Student’s *t* test.

Likewise, a primary mouse model was established with liver-specific overexpression of SLC13A2 using AAV8 under the control of the TBG promoter. After 2 weeks of AAV injection, the mice were injected with oncogene plasmids via the HTVi method and sacrificed after 9 more weeks, which is a longer period than that of the LKO experiments, to better demonstrate tumor suppression by SLC13A2 (Figure 4A). The overexpression efficiency of SLC13A2 was verified at the mRNA and protein levels (Figure 4B). The number of surface tumor nodules and liver/body weight ratio were significantly lower in the SLC13A2-overexpressing group than in the control group, while no significant differences were observed in body weight or blood glucose between the two groups (Figure 4C-D). The tumor burden severity was significantly different, as was the percentage of tumor incidence. The maximal tumor size in the SLC13A2 overexpression group was significantly reduced compared to that in the GFP group (Figure 4E). The enzymatic activity of ALT and AST was consistently inhibited in the blood and liver tissues harvested from mice with liver-specific overexpression of SLC13A2 (Figure 4F). Histological staining revealed that hepatocytes were smaller and more separated in the livers of the SLC13A2-overexpressing group than in those of the control group, as indicated by a decreased nucleo-plasmic ratio (Figure 4G). Consistent with the decrease in tumor growth, immunohistochemical staining of Ki67 also revealed 8.0% nuclear positivity in the control group and approximately 2.8% nuclear positivity in the SLC13A2 overexpression group, demonstrating a significant decrease in cell proliferation induced by SLC13A2 *in vivo* (Figure 4H). Similarly, the RNA expression of cell proliferation-associated genes, such as *Ccnb1*, *Foxm1, Fgf21* and *Mcm2,* decreased in liver tissues overexpressing SLC13A2 *in vivo* (Figure 4I). Moreover, the expression of genes involved in metabolism, such as *Hk2* and *Mdh2,* also significantly decreased with the overexpression of SLC13A2 (Figure 4I). These findings indicated a favorable metabolic status for HCC progression induced by SLC13A2.

**Figure 4.**
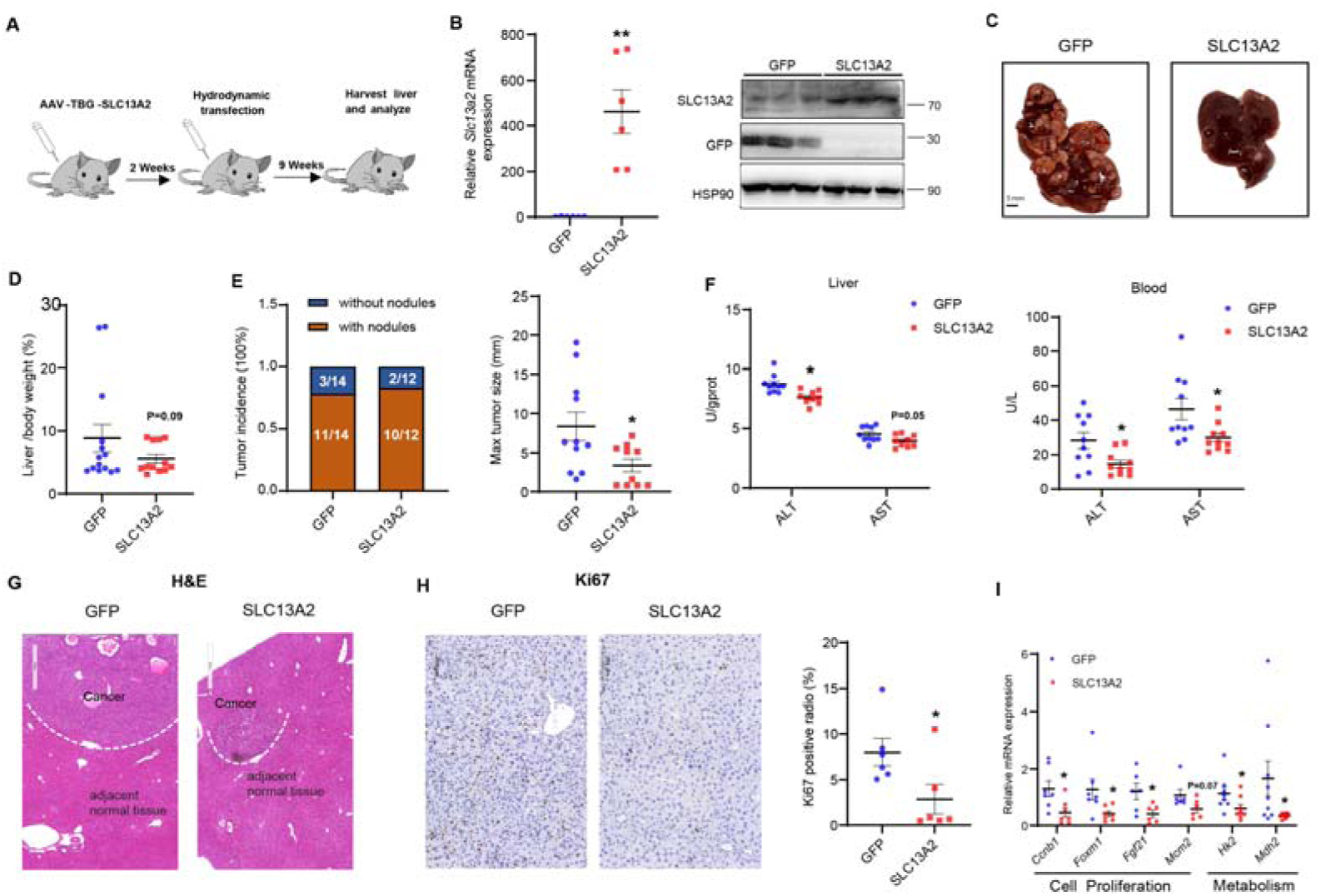
Liver-specific overexpression of SLC13A2 inhibits HCC tumor progression. **A.** Schematic diagram of the liver-specific overexpression of SLC13A2 and study design. AAV-TBG-GFP (GFP) and AAV-TBG-SLC13A2 (SLC13A2) viruses (2 × 10^11^ VP/mouse) were injected into wild-type mice via the tail vein. The transduced mice were subjected to the HTVi model by injecting SB in combination with oncogene plasmids (c-MET: ΔN90-β-catenin: SB = 20 μg:20 μg:5 μg). **B.** qPCR analysis (left, n=6 per group) and immunoblots (right) showing SLC13A2 overexpression efficiency in mouse HCC tissues. **C.** Representative images showing liver morphology upon dissection. Scale bar: 5 mm. **D.** Liver/body weight ratio (n=14 for the GFP group and n=12 for the SLC13A2 OE group). **E.** Tumor incidence showing the number of mice with or without nodules (left) and the maximal tumor size (right, n=12 for the GFP group and n=10 for the SLC13A2 OE group) in the two groups. **F.** Liver and serum ALT/AST activity. **G.** Representative images of H&E staining. **H.** Immunohistochemical staining for Ki67. Ratio of Ki67-positive cells in the two groups (right, n=6 per group). **I.** qPCR analysis of liver tissues showing gene transcription for cell proliferation and metabolism. The data are presented as the means ± SEMs. *P < 0.05, **P < 0.01, SLC13A2-OE vs GFP group, two-tailed unpaired Student’s *t* test.

### SLC13A2 inhibits HCC cell proliferation by depleting TCA cycle intermediates for macromolecular biosynthesis and suppressing oxidative respiration

Next, a mouse HCC cell line, Hepa1-6, and a human HCC cell line, HepG2, were chosen to confirm the capacity of SLC13A2 to prevent the proliferation of HCC cells. To further investigate the effect of SLC13A2 on the proliferation of HCC cells *in vitro*, we used RNA interference (KD) to knock down SLC13A2 and plasmid/adenovirus transfection to induce SLC13A2 expression (OE) in HCC cells. Notably, RNA interference transduction significantly reduced the protein expression of SLC13A2 in Hepa1-6 cells (Figure 5A). Strikingly, growth curve and colony formation experiments revealed that the colony formation ability and viability of Hepa1-6 cells were significantly increased with SLC13A2 knockdown (Figure 5B-C). Transfection of Hepa1-6 or HepG2 cells with plasmids or adenoviruses increased the protein expression of SLC13A2 (Figure 5A). Growth curve and colony formation assays revealed that SLC13A2 overexpression significantly decreased the viability and clonogenicity of Hepa1-6 and HepG2 cells (Figure 5B-C). To further confirm the effect of SLC13A2 on cell proliferation, an EdU incorporation assay was applied. The ratio of EdU-positive cells was significantly greater in SLC13A2-KD cells than in SLC13A2-OE cells (Figure 5D). Consistently, a cell cycle distribution assay showed that SLC13A2 knockdown decreased the proportion of cells in S phase, while SLC13A2 overexpression increased S phase arrest (Figure 5E).

**Figure 5.**
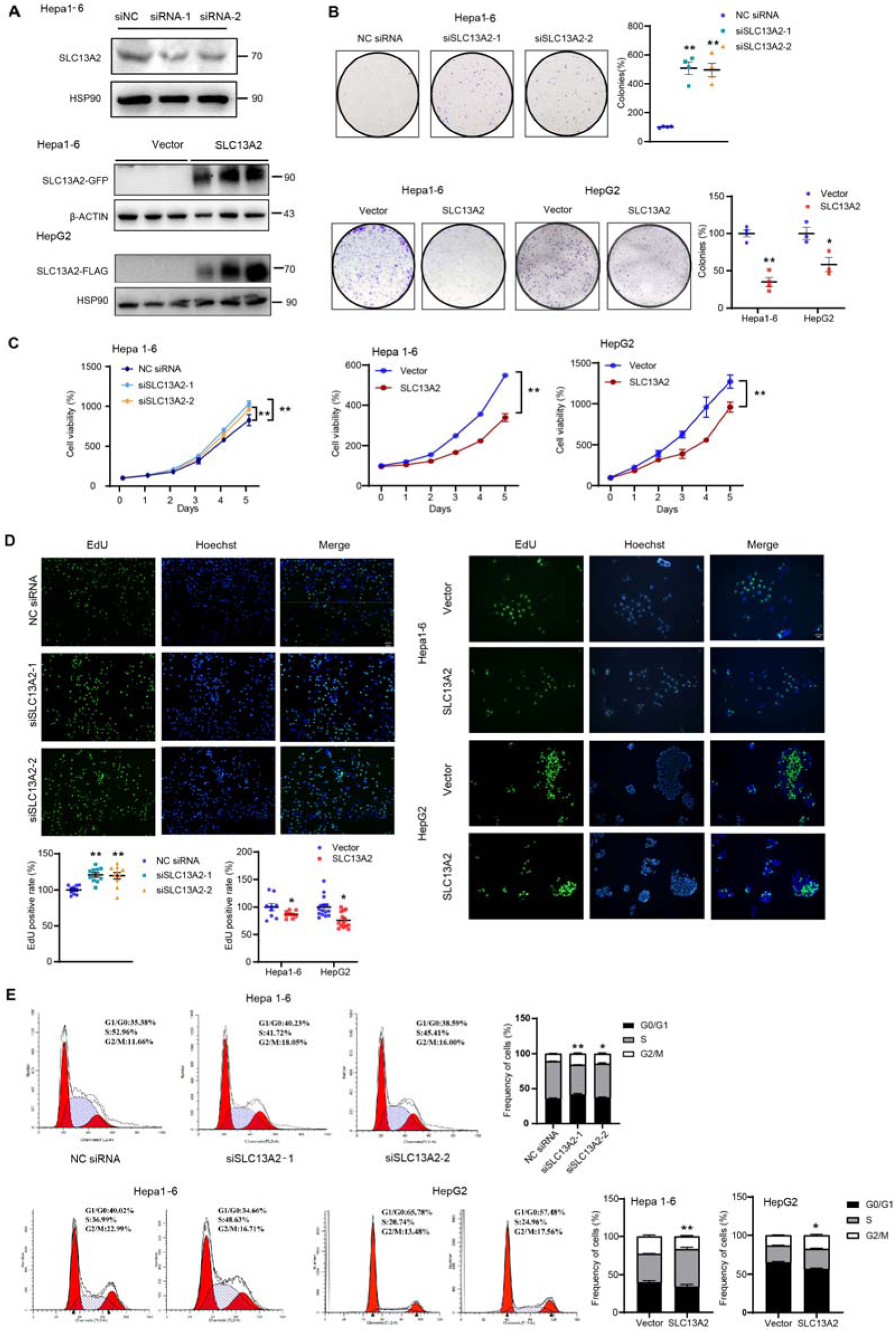
SLC13A2 inhibits the proliferation of HCC cells *in vitro*. **A.** Knockdown and overexpression efficiency *in vitro.* The protein level of SLC13A2 in Hepa1-6 cells was evaluated after transfection with NC/vector or SLC13A2 siRNA/expression plasmids/adenovirus for 48 h. **B.** Colony formation assay showing the clonogenic potential of HCC Hepa1-6 cells after treatment with SLC13A2 KD (siSLC13A2, top), Hepa1-6 cells and HepG2 cells with SLC13A2 OE (bottom) and culture for 7 or 14 days. The number of colonies was counted. **C.** The growth of HCC cells with SLC13A2 knockdown (left) or overexpression (right) was analyzed by CCK8 assays at 0, 24, 48, 72, 96, and 120 h after transfection. **P < 0.01, two-way ANOVA, followed by Bonferroni multiple comparisons. **D.** EdU incorporation was detected in Hepa1-6 cells with SLC13A2 KD (left) or in Hepa1-6 and HepG2 cells with OE (right) for 72 h, followed by incubation with EdU (20 μM) for 3-4 h (scale bar: 200 μm or 50 μm). EdU-positive cells were visualized under a fluorescence microscope, and the percentages are shown below. **E.** Propidium iodide (PI) staining and flow cytometry analysis showing cell cycle arrest in Hepa1-6 cells after transfection with SLC13A2 siRNAs or control siRNA (NC) for 48 h (top) or in Hepa1-6 and HepG2 cells after overexpression of SLC13A2 for 72 h (bottom). Bars represent the means ± SEMs (n=3 or 4 independent experiments). *P < 0.05 and **P < 0.01, two-tailed unpaired Student’s *t* test (B, D-E).

Since classical substrates of SLC13A2 are TCA cycle intermediates that control cellular function and cell fate (3), we conducted UPLC-QTOF-MS-based untargeted metabolomics to investigate the effect of SLC13A2 overexpression on metabolism in HCC Hepa1-6 cells. The levels of intermediates of the TCA cycle, such as citrate, a-KG, malate and aconitate; nonessential amino acids, such as glutamine and aspartate; and metabolites involved in nucleotide metabolism, such as cytidine and orotidine, decreased after SLC13A2 overexpression (Figure 6A). The differentially abundant metabolites were largely enriched in alanine-aspartate-glutamate metabolism, D-glutamine and D-glutamate metabolism and the aminoacyl-tRNA biosynthesis pathway (Figure 6B). Then, the effects of SLC13A2 on oxidative respiration were detected using Seahorse analysis. SLC13A2 overexpression impaired basal and maximal respiratory capacity (Figure 6C). Consistently, ATP production decreased with SLC13A2 overexpression and increased with SLC13A2 knockdown (Figure 6D). To detect the mitochondrial transmembrane potential, the cells were stained with the sensitive fluorescent probe JC-1. Mitochondrial depolarization was induced by SLC13A2, as the ratio of aggregates to monomers significantly decreased with SLC13A2 overexpression and increased with SLC13A2 knockdown (Figure 6E). This finding suggested that SLC13A2 decreases cellular mitochondrial capability by inhibiting oxidative phosphorylation and ATP synthesis and increasing mitochondrial permeability.

**Figure 6.**
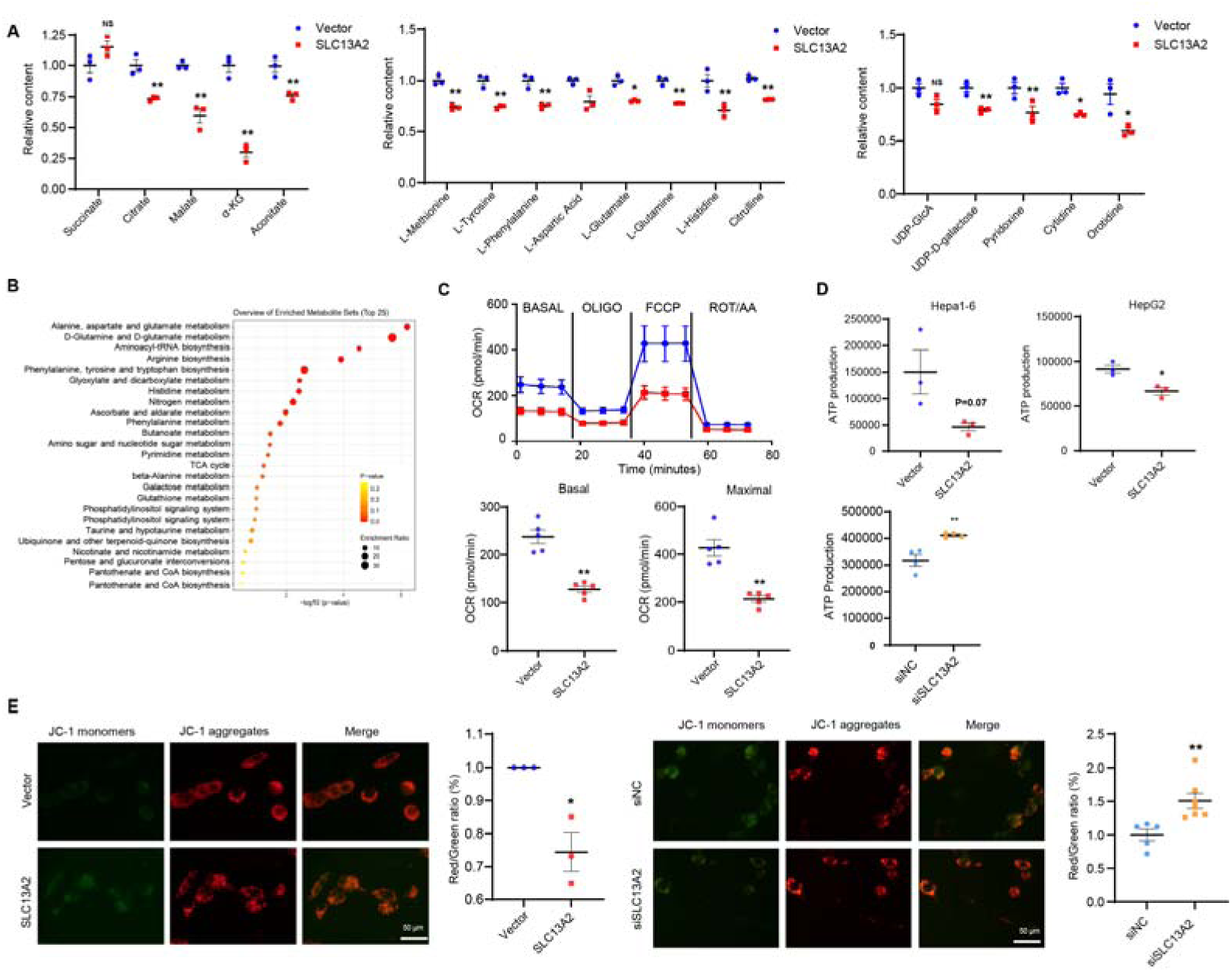
SLC13A2 restricts the TCA cycle and inhibits mitochondrial respiration. **A-B.** SLC13A2 was overexpressed in Hepa1-6 cells, which were subsequently cultured for an additional 60 h, after which the metabolites were analyzed via untargeted metabolomics. (**A)** Relative content analysis of TCA cycle substrates (left), amino acid metabolites (middle) and nucleotide metabolites (right). (**B)** Metabolite set enrichment analysis (MSEA) of the metabolic pathways (n = 3). adj. P < 0.05 was considered to indicate statistical significance. **C.** Hepa1-6 cells were cultured for 24 h after transfection with the vector or SLC13A2 overexpression plasmid. After the cells were digested, 8,000 cells/well were seeded for the Seahorse XF assay. Dashed vertical lines indicate the subsequent addition of the ATPase inhibitor oligomycin (OligO: 1.5 μM), the uncoupling reagent (FCCP: 1.0 μM) and the inhibitors of the electron transport chain rotenone/antimycin A (ROT/AA: 0.5 μM). The oxygen consumption rate (OCR) under basal conditions and after treatment with the uncoupler FCCP (maximal respiration) were analyzed. **D.** HepG2 and Hepa1-6 cells were cultured for 72 h after transfection to detect ATP production. **E.** Confocal images of JC-1 fluorescence. The graph shows the ratio of aggregates (red) to monomers (green), indicating changes in the mitochondrial membrane potential. The data are presented as the means ± SEMs (n=3 independent experiments). *P < 0.05 and **P < 0.01, two-tailed unpaired Student’s *t* test (A, C-E).

### Citrate-imported SLC13A2 acetylates and thus degrades PKM2, resulting in TCA cycle restriction to suppress HCC cell proliferation

The mechanisms by which SLC13A2, a well-known citrate inward transporter, restricts the TCA cycle to suppress HCC tumor growth are intriguing. Therefore, we used metabolic flux analysis to better understand the effects of SLC13A2 on glycolytic flux via the application of U-^13^C_6_-glucose at the indicated time points. Surprisingly, after SLC13A2 was overexpressed, the levels of the upstream glycolysis intermediates G-6-P, GAP/DHAP, 3-PG, and PEP significantly increased, while the levels of the downstream metabolite pyruvate significantly decreased with SLC13A2 overexpression (Figure 7A). Furthermore, targeted metabolomic analysis confirmed that the intracellular pyruvate content decreased significantly with SLC13A2 overexpression, which was consistent with the restoration of cell proliferation by an additional supply of pyruvate (500 µM) in Hapa1-6 cells. (Figure 7B). These findings suggested that SLC13A2 controls the entrance of the TCA cycle, which was consistent with its ability to suppress tumor growth and attenuate cell proliferation.

**Figure 7.**
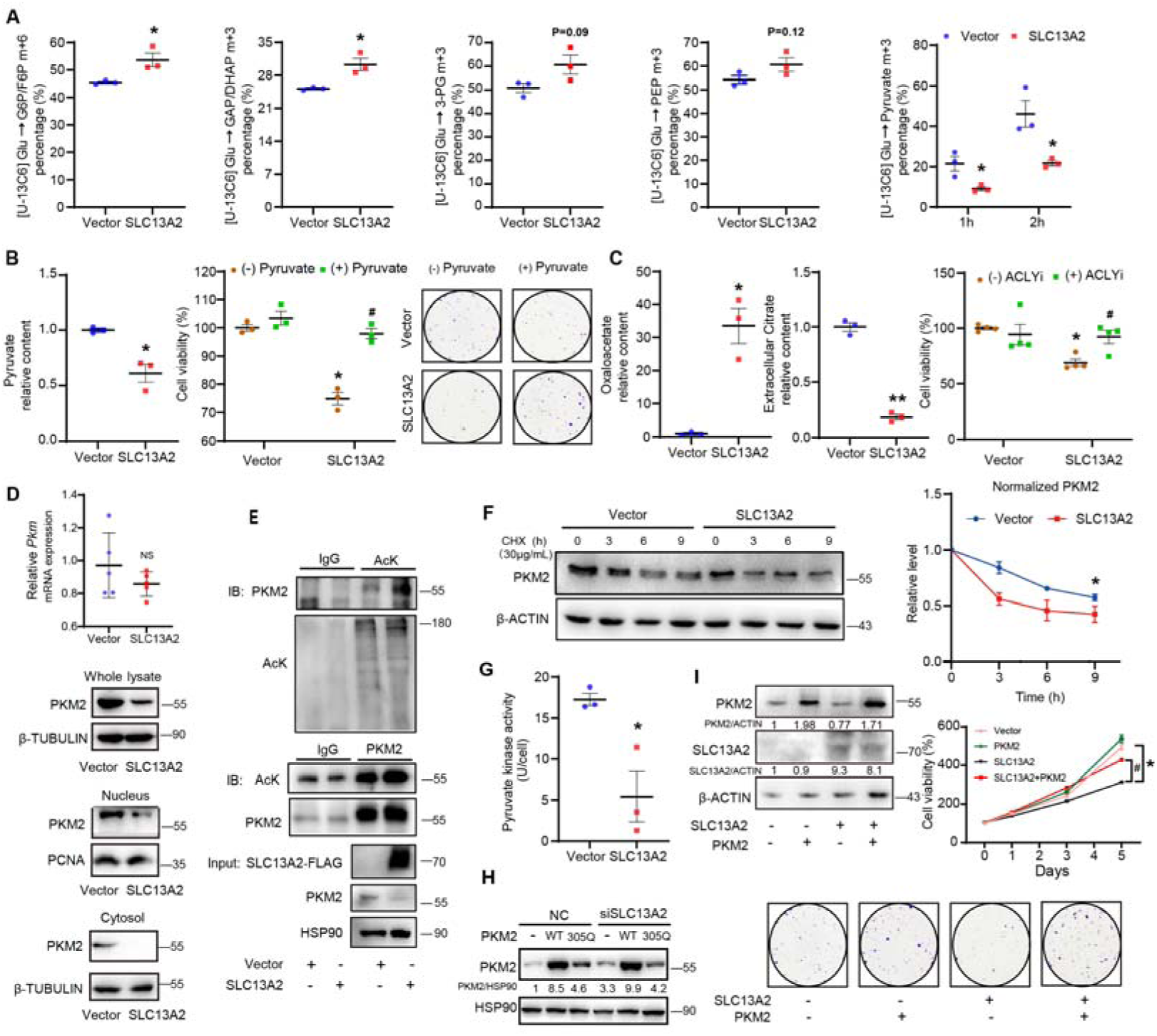
SLC13A2 suppresses tumor growth through the acetylation and degradation of PKM2 by imported citrate. **A.** Hepa1-6 cells were transfected with vector or SLC13A2-OE adenovirus and cultured for an additional 60 h. U-^13^C_6_-glucose was then added, and the cells were incubated for 1 h (A-D) or the indicated period (E). The metabolites were analyzed using an untargeted metabolomics method. **B.** Hepa1-6 cells were transfected with vector or SLC13A2-OE adenovirus and cultured for 60 h. The relative content of pyruvate was analyzed using targeted metabolomics (left). The viability of Hepa1-6 cells with or without SLC13A2 overexpression was measured after treatment with pyruvate for 72 hours (middle). Cell colony formation assay showing the clonogenic potential of HCC cells after treatment with pyruvate for 7 days in the presence or absence of SLC13A2 overexpression (right). *P < 0.05 vs. vector without pyruvate, ^#^P < 0.05 vs. SLC13A2 without pyruvate. **C.** Hepa1-6 cells were transfected with vector or SLC13A2-OE adenovirus and cultured for 60 h. The relative levels of intracellular oxaloacetate (left) and extracellular citrate (middle) were analyzed via targeted metabolomics. Hepa1-6 cell viability was measured after treatment with an ACLY inhibitor for 72 hours in the presence or absence of SLC13A2 overexpression (right). *P < 0.05 vs. vector without the ACLY inhibitor, ^#^P < 0.05 vs. SLC13A2 without the ACLY inhibitor. **D.** PKM2 RNA or protein levels in whole-cell lysates, nuclei and cytosol of Hepa1-6 cells after transfection with the vector or SLC13A2 and culture for 60 h. **E.** Co-IP assay showing the acetylation of PKM2 in Hepa1-6 cells with or without SLC13A2 overexpression. IgG served as a negative control. **F.** Cycloheximide (CHX) chase assay showing the effect of SLC13A2 on the half-life of the PKM2 protein in Hepa1-6 cells by immunoblotting (left) and densitometry analysis (right). **G.** Pyruvate kinase activity in Hepa1-6 cells after transfection for 60 h. **H.** Hepa1-6 cells were transfected with vector or SLC13A2-OE adenovirus in the presence or absence of WT or mutant PKM2. Immunoblots showing the PKM2 protein level after culture for 60 h. **I.** Immunoblot analysis showing the SLC13A2 and/or PKM2 overexpression efficiency in Hepa1-6 cells after transfection. The growth curves of Hepa1-6 cells overexpressing SLC13A2 and/or PKM2 were analyzed by CCK8 assays at 0, 24, 72 and 120 h after transfection. Cell colony formation assay showing the clonogenic potential of HCC cells after transfection with SLC13A2 and/or PKM2 and culture for 7 days. *P < 0.05 vs. Vector, ^#^P<0.05 vs. SLC13A2. The data are presented as the means ± SEMs (n=3 or 4 independent experiments). *P < 0.05, two-tailed unpaired Student’s *t* test (A, B left, C left and middle, G); *P < 0.05, two-way ANOVA, followed by Bonferroni multiple comparisons (B middle, C right, F, I).

Since citrate is one of the major substrates of SLC13A2 with the highest affinity (20), we hypothesized that the citrate imported by SLC13A2 was catalyzed by ACLY to produce oxaloacetate and acetyl-CoA. We used targeted metabolomic analysis to detect intracellular oxaloacetate and found that it was significantly increased by SLC13A2 overexpression, while extracellular citrate was significantly decreased (Figure 7C, left). These results indicated that SLC13A2 imported extracellular citrate, which produced oxaloacetate. Cell viability assays also revealed that treatment with an ACLY inhibitor (50 µM) significantly reversed the SLC13A2-mediated inhibition of cell growth (Figure 7C, right). As a major enzyme that catalyzes the transition of PEP to pyruvate in HCC cells (21), PKM2 was therefore suggested to be regulated by SLC13A2. We thus hypothesized that acetyl-CoA released by citrate catalysis may act as a precursor for the acetylation of PKM2, which leads to its protein degradation (22). To evaluate the protein distribution of PKM2 after SLC13A2 overexpression, we prepared isolated subcellular fractions from the nucleus and cytosol and found that SLC13A2 significantly decreased PKM2 protein levels in the cytosol and nucleus (Figure 7D below). While the PKM2 protein was decreased by SLC13A2, the RNA level was unchanged (Figure 7D up), which also indicated that posttranslational modification resulted in a decrease in the PKM2 protein. Therefore, we performed coimmunoprecipitation (co-IP) experiments in SLC13A2-overexpressing Hepa1-6 cells. The acetylation of PKM2 increased significantly with SLC13A2 overexpression (Figure 7E). We further used a CHX-chase assay to examine whether SLC13A2 accelerated the turnover of the PKM2 protein. The data revealed that protein degradation was the primary pathway involved in the SLC13A2-mediated decrease in the PKM2 protein level (Figure 7F). Consistently, pyruvate kinase activity was strongly attenuated by SLC13A2 overexpression (Figure 7G). Since K305 has been reported to be acetylated for PKM2 protein degradation(22), we generated a PKM2-K305Q acetylation-mimic mutant and transfected it into HCC cells with SLC13A2 knockdown. The results confirmed that the level of the PKM2 305Q mutant protein significantly decreased compared with that of the WT protein. The knockdown of SLC13A2 increased the levels of both the endogenous protein and the transfected WT PKM2 protein but did not increase the levels of the K305Q mutant protein (Figure 7H). Furthermore, PKM2 overexpression reversed the inhibitory effect of SLC13A2 on HCC cell proliferation, as confirmed by growth curve and colony formation analyses (Figure 7I). SLC13A2 suppresses HCC tumor growth in a PKM2-dependent manner.

### SLC13A2 remodels gene transcription for proliferation and metabolism, which restrains tumor growth

To delineate the mechanisms of SLC13A2 in tumor suppression, we evaluated the transcriptional regulation induced by SLC13A2 using transcriptomic RNA-seq analysis of total liver RNA from the GFP and SLC13A2-OE groups in the HTVi model (Figure 4A). We identified a total of 2205 hepatic DEGs in SLC13A2-OE mice (FDR<0.05, fold change>2, adj P>0.05) (Figure 8A). Gene Ontology analysis revealed that the downregulated genes were enriched in cell proliferation and nuclear division, and the upregulated genes were enriched mainly in small molecule catabolic pathways and nutrient metabolism pathways (Figure 8B). Gene set enrichment analysis also revealed that 102 gene sets were significantly differentially regulated by SLC13A2. The top two gene sets of organic acid catabolic process and amino acid catabolic process were downregulated in the SLC13A2-OE group (Figure 8C). It has been reported that PKM2 not only acts as a PK enzyme but is also is located in the nucleus to promote gene transcription as a transcription factor or cotransporter by physically interacting with other transcription factors, including HIF-1α, the NF-κB subunit p65/RELA, STAT3 and ATF2 (23–25). In addition, nuclear PKM2 also induces the expression or regulates the protein stability of transcription factors such as EGR1 and RUNX1 (26). By analyzing transcription factor binding motif enrichment, relevant transcription factors, including RELA, EGR1, HNF4a, STAT3 and RUNX1, which are highly associated with PKM2 nuclear functions, were predicted to be involved in the regulation of gene transcription by SLC13A2 during HCC tumorigenesis (Figure 8D).

**Figure 8.**
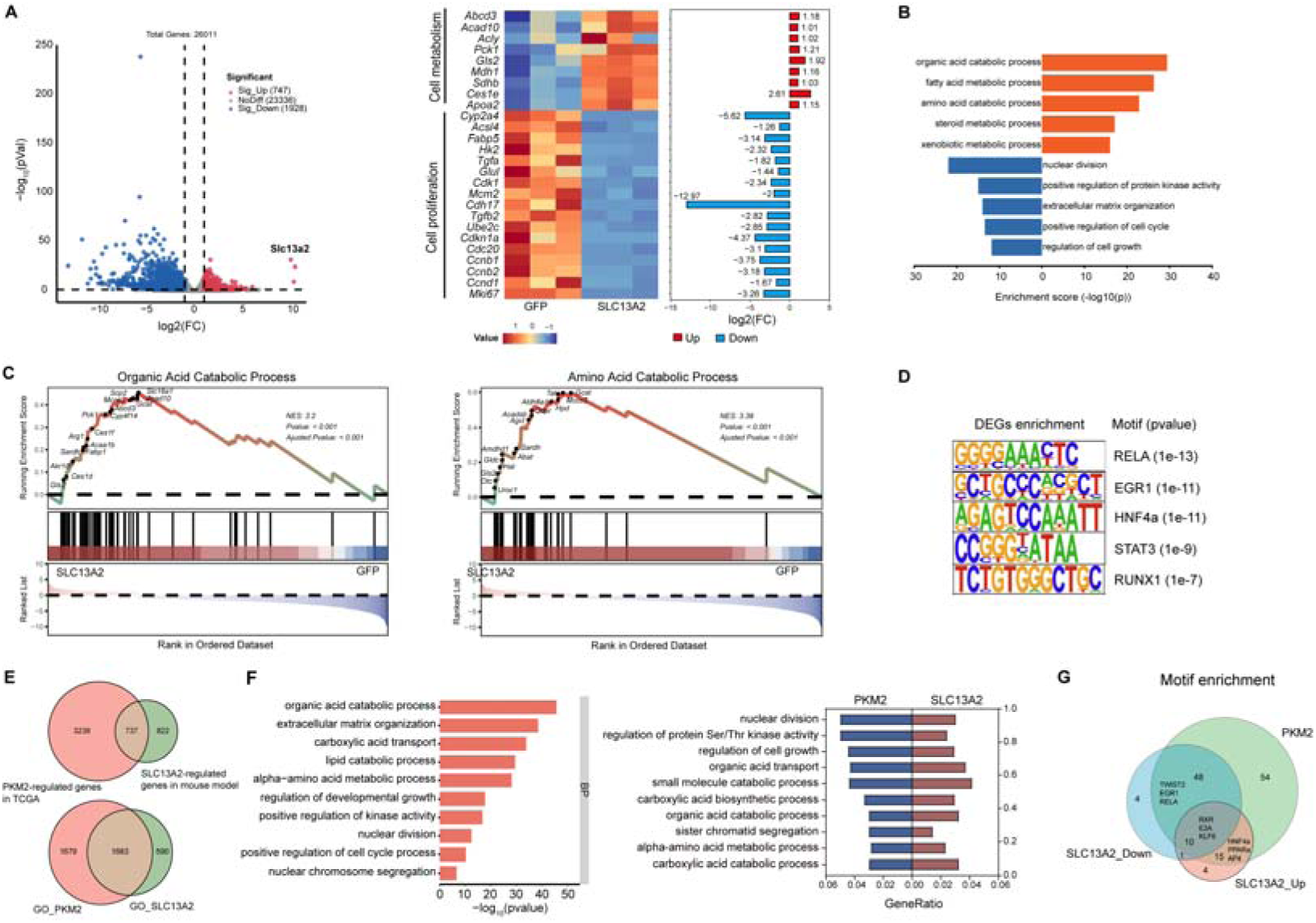
SLC13A2 suppresses tumor growth by reprograming PKM2-dependent gene transcription for cell proliferation and metabolism. **A.** Volcano plot of hepatic genes upregulated or downregulated in SLC13A2-OE mice (left). Representative genes involved in cell proliferation and metabolism are shown (right). **B.** Enrichment of biological processes in the two clusters is indicated for SLC13A2-OE mice. **C.** Gene set enrichment analysis of all DEGs in SLC13A2-OE mice, showing the top two DEGs. **D.** Motif enrichment analysis of predicted transcription factors in SLC13A2-OE mice. **E.** The Venn diagrams illustrates the conservative DEGs shared between PKM2-regulated genes in the TCGA HCC level 3 clinical patient cohort and the SLC13A2-regulated genes in HCC mouse model (top), and the enrichment of biological processes shared by the regulatory gene clusters regulated by PKM2 and SLC13A2 (bottom). **F.** Gene ontology enrichment by the PKM2-regulated gene cluster (left). The representative enriched biological processes shared by the regulatory gene clusters of PKM2 and those of SLC13A2 (right). **G.** Motif enrichment showing representative predicted transcription factors shared among the clusters of PKM2-regulated, SLC13A2-upregulated and SLC13A2-downregulated genes.

Furthermore, we analyzed the RNA-seq data of 371 primary HCC patients in level 3 from the TCGA clinical dataset and evaluated the differential genes in the PKM2-high group compared with PKM2-low expression group (27). These differential genes reflected the regulatory effects of PKM2 on gene expression. We identified 3975 differentially expressed genes that were conserved between human and mouse, with a coverage of approximately 50% SLC13A2-regulated genes (Figure 8E top). Gene ontology enrichment showed similar gene expression patterns between the regulation of PKM2 and the induction by the overexpression of SLC13A2 (Figure 8E bottom and 8F). By performing transcription factor motif enrichment analysis, we observed that transcription factor-binding sites predicted for the genes whose transcription was differentially regulated by SLC13A2 were also highly enriched for the genes whose expression was differentially regulated by PKM2 in human HCC patient datasets. The shared motifs enriched for the expression of both SLC13A2- and PKM2-regulated genes corresponded to predicted binding sites for transcription factors, including RELA, EGR1, HNF4a, RXR and PPARa (Figure 8G). The PKM2-regulated genes covered about 90% of the predicted transcription factors by SLC13A2, indicating the dependence of SLC13A2-regulated gene expression on nuclear PKM2. The data establish a novel model of PKM2 protein degradation through the SLC13A2-mediated inward transport of citrate, which acts as a signaling metabolite that suppresses HCC progression (Graphic abstract).

## Discussion

Since the ‘Warburg effect’ was proposed by Otto Warburg during the 1920s, an increasing number of scholars have focused on the role of aerobic glycolysis in tumors, but the role of the TCA cycle, not to mention the TCA cycle substrates, is still largely obscure (28). In the present study, we found that SLC13A2, an SLC transporter that imports citrate, is downregulated in human and mouse HCC cells.

Citrate produces acetyl-CoA for the acetylation and degradation of PKM2, thus depleting TCA cycle intermediates and decreasing macromolecule biosynthesis and gene transcription for HCC tumor growth (schematic diagram shown in the graphic abstract). As deregulated metabolism is considered a key hallmark of HCC progression, our findings suggest a novel transporter with potential for the design of drugs or strategies for HCC treatment.

Citrate is a substance involved in the TCA cycle and de novo fatty acid/cholesterol synthesis and plays a key role in cell survival. Deciphering the consequences of citrate metabolism is important for dissecting the etiology of HCC. The classical plasma membrane citrate transporters SLC13A2, SLC13A3, SLC13A5 and mitochondrial SLC25A1 have nonnegligible effects on the metabolism of citrate.

SLC13A family transporters, which can be divided into two functionally distinct groups, mediate the movement of Na+-coupled anion substrates across the membrane (29). The group of Na^+^-di- and tri-carboxylate cotransporters carrying TCA cycle intermediates includes SLC13A2, SLC13A3 and SLC13A5, among which the first is exclusively downregulated in HCC liver tissues and cells (Figure 1B, 1F and 2A-C, 2F). As a previously defined transporter expressed in the apical membrane of renal proximal tubular cells and small intestine cells, SLC13A2 is considered a primary protein involved in citrate reabsorption and is thus important for acidLbase homeostasis (30). In renal tubules, SLC13A2 integrates the movement of three sodium ions with dicarboxylate and is triggered by a low intracellular sodium concentration as well as intracellular electronegativity. SLC13A2 has high specificity for dicarboxylates with different affinities for substrates (0.35 mM km for succinate and 0.6 mM km for citrate) (31). Although SLC13A2 has a greater affinity for succinate than citrate, as previously described, the latter is a much more abundant metabolite in HCC cells than the former (the content ratio is more than 2800-fold) (Figure S1C). There is also less intracellular a-KG than citrate (approximately 14-fold). To date, few studies have focused on the role of SLC13A2 in addition to the kidney, especially the liver. In contrast to the general belief that SLC13A2 is expressed at very low levels in the liver, we found that SLC13A2 mRNA is normally expressed in healthy liver tissue (Table 1), indicating its role in the physiological homeostasis of liver function. Although we did not observe any obvious phenotype with hepatocyte-specific overexpression or knockout of SLC13A2 (Figure S2, S3) under physiological conditions, it possibly contributes to tuning the metabolic status of hepatocytes for the maintenance of liver homeostasis. We speculate that a robust phenotype might be shown with metabolic stresses such as HFD feeding. The role of SLC13A2 in the regulation of liver metabolism still deserves further elucidation.

As another member of the SLC13 family, SLC13A5 has gained increasing attention because of its high expression in the liver. SLC13A5 reportedly regulates hepatic glucose and fatty acid metabolism by transporting citrate from the extracellular to the intracellular compartment. Knockout of SLC13A5 prevents high-fat diet-induced insulin resistance by attenuating hepatic gluconeogenesis and lipogenesis (32). RNAi-mediated silencing of SLC13A5 expression in two human HCC cell lines, HepG2 and Huh7, profoundly suppressed cell proliferation and colony formation and induced cell cycle arrest through increased expression of the cyclin-dependent kinase inhibitor p21 and decreased expression of cyclin B1 (33). However, the mRNA expression of both *Slc13a3* and *Slc13a5,* two other members with similar functions in the family, did not significantly or consistently change in HCC liver tissues (Figure 1B, 1F and 2B). The extracellular import of citrate induced by SLC13A2 represents a new mechanism for regulating HCC metabolism that is distinct from that of SLC13A5.

SLC25 family members are considered important transporters of the inner mitochondrial membrane and are involved in numerous metabolic pathways (34). SLC25A1 (also known as mitochondrial citrate transport protein) is a mitochondrial carrier that transports citrate or isocitrate into the cytosol in exchange for malate. Increased transcription and increased activity of SLC25A1 have been revealed as hallmarks of several metabolic disorders and several cancer types (35). It has been reported that SLC25A1 is highly expressed in the metastatic lymph nodes of non-small cell lung cancer patients. Lung cancer stem cells rely on SLC25A1 to maintain the mitochondrial citrate pool and redox balance, resulting in the stimulation of oxidative phosphorylation for energy production and cell survival (36). Studies have shown that SLC25A1 promotes the entry of cytosolic citrate into mitochondria through reverse import activity, revealing the obscure role of SLC25A1 in tumorigenesis (36). Anchorage-independent tumor growth requires changes in the metabolism of citrate, which is imported into mitochondria by SLC25A1, to inhibit mitochondrial oxidative stress (37). In addition, SLC25A1 plays an important role in lipid metabolism by exporting citrate from mitochondria, and pharmacological inhibition of SLC25A1 prevents NASH pathogenesis (38). As another key dicarboxylate carrier, SLC25A10 transports malate and succinate out of mitochondria in exchange for phosphate, sulfate and thiosulfate. The expression of SLC25A10 has been linked to cancer, but further understanding of its role in mitochondrial function and TCA cycle intermediates is lacking (39, 40). A recent study revealed that SLC25A10 is predominantly expressed in white adipose tissue and is responsible for succinate transport from mitochondria to the cytosol. The exported succinate therefore acts on its receptor SUCNR1 to inhibit adipocyte lipolysis and protect against liver lipotoxicity upon HFD feeding (41). Both SLC25A1 and SLC25A10 transport substrates by electroneutral exchange (42), which is distinct from the scenario induced by SLC13A2. Downregulation of SLC13A2 in HCC liver tissues and cells maintains the mitochondrial membrane potential across the inner membrane (Figure 6E), which ensures the functional operation of the TCA cycle for oxidative phosphorylation and ATP production in HCC cells (Figure 6C-D).

Apart from playing a central role in mitochondrial metabolism and respiration, citrate also plays fundamental roles in the cytosol by serving as a metabolic substrate, an allosteric enzymatic modulator and a source of acetyl-CoA as an epigenetic modifier and precursor for lipogenesis (43). The activities of enzymes involved in glycolysis, lipogenesis and gluconeogenesis are regulated by binding to citrate, for example, the activation of 1,6-bisphosphatase (FBP1) and acetyl-CoA carboxylase alpha (ACACA) and the inhibition of phosphofructokinase (PFK) (44, 45). Our results reveal a novel mechanism for subtly tuning the intracellular concentration of citrate for HCC cell growth. SLC13A2 imports citrate, leading to the inhibition of glycolysis through the acetylation and degradation of PKM2 (Figure 8D-F). This finding echoes previous findings that the intracellular concentration of citrate remains low in cancer cells to sustain tumor growth (46). Intriguingly, SLC13A2 also gates the entrance to the TCA cycle and decreases its intermediates, including citrate (Figure 6A left), for cell growth. This indicates that the depletion of TCA cycle intermediates is the eventual consequence of SLC13A2, which ultimately suppresses cell growth. The present findings provide further evidence that citrate serves as a metabolic signal to regulate cell growth in addition to serving as a metabolite in the TCA cycle to produce energy.

Aerobic glycolysis is found in HCC and is a hallmark of liver cancer and is responsible for the regulation of HCC proliferation, immune evasion, invasion, metastasis, angiogenesis and drug resistance. Pyruvate kinase (PK) is an enzyme that catalyzes the conversion of phosphoenolpyruvate and ADP to pyruvate and ATP in glycolysis and plays a role in regulating cellular metabolism. Unlike other PKs, PKM2 is highly upregulated in cancer cells and is associated with a poor prognosis (47). The activity of PKM2 can also be regulated by posttranslational modifications, including acetylation, oxidation, phosphorylation, prolyl hydroxylation, and SUMOylation. These modifications have been shown to modulate PKM2 activity, degradation, oligomerization, and subcellular localization. PKM2 is directly phosphorylated on tyrosine residues, including Y105, allowing phosphorylated PKM2 to play a role in inactivating other PKM2 tetramers by stimulating FBP release (48). Acetylation of PKM2 K305 reduces pyruvate kinase activity by reducing its affinity for PEP and triggering degradation of the enzyme itself, and acetylation of PKM2 K433 disrupts FBP binding and reduces enzyme activity (22, 49). A decrease in PK activity suppresses glycolysis and gates the access of nutrients to the TCA cycle (Figure 7G). These effects restrain the TCA cycle for the endogenous production of TCA intermediates, including citrate (Figure 6A, left), and lead to an insufficient supply of building blocks for the synthesis of macromolecules, including amino acids and nucleotides (Figure 6A, right), which are essential for cancer cell growth (4). Our study revealed that SLC13A2 inhibits HCC cell proliferation by inducing the acetylation and degradation of PKM2 (Figure 7D-F).

Our data indicate that SLC13A2 is a potential drug target for HCC treatment. Recent RNA-seq data analysis revealed that several anticancer drugs, including nilotinib and imatinib (GSE19567), elevated the transcription of SLC13A2. It has been reported that the transcription factor HNF4a is involved in the downregulated expression of SLC13A2, suggesting that saturated fatty acid and inflammatory signals are factors that inhibit SLC13A2 gene transcription (GSE210842). Similarly, a HFD potently decreases the expression of SLC13A2 (GSE68360). A deep understanding of the regulatory mechanisms of SLC13A2 expression during HCC tumorigenesis will provide insightful suggestions for drug discovery strategies targeting this transporter.

To date, few studies have linked SLC13A2 with cancer, and no investigations have investigated its roles and mechanisms in cancer pathogenesis. According to the database of the Human Protein Atlas (www.proteinatlas.org) (50), the expression of SLC13A2 has low cancer specificity. A high expression level of SLC13A2 is positively associated with a better overall survival rate in patients with renal cancer, suggesting its role in improving the prognosis of cancer patients. Transcriptome analysis of primary colorectal cancer tissues from patients revealed that SLC13A2 is a representative differentially expressed gene that is responsible for liver metastasis (51). Another study conducted a genome-wide association analysis with familial HBV-related HCC in comparison with non-HCC controls with chronic HBV to identify susceptibility loci (52). The results revealed a single-nucleotide polymorphism cluster located at the 3’ end of SLC13A2, which is among the genes most strongly associated with familial HBV-associated HCC (52). This finding is in accordance with our findings suggesting that SLC13A2 is a hereditary genetic factor related to the pathogenesis of HCC.

## Statement of significance

SLC13A2 is a downregulated SLC transporter for TCA cycle intermediates and plays a protective role during HCC progression. SLC13A2-transported citrate acts as a precursor to acetylate the glycolytic enzyme PKM2 and leads to its degradation. Decreased nuclear PKM2 remodels gene transcription for cell proliferation and metabolism, which are required for tumor growth.

## Author contributions

Q.M., S.L., C.H., L.C., T.Z., H.D. and S.C. performed the research; X.J. conceived the study; X.J. and H.H. designed the research study; X.J. and H.H. contributed essential reagents or tools; X.J., C.H., A.D. and Y.W. analyzed the RNA-seq datasets; Y.H. collected the clinical samples; Q.M., L.C. Y.S. and X.J. analyzed the data; and X.J., Q.M., and L.C. wrote the paper. All the authors have read and approved the final manuscript.

## Acknowledgments

This study was supported by the National Natural Science Foundation of China (No. 82070883 and 82273982 to J.X.), the National Natural Science Foundation of Jiangsu Province (No. BK20221525 to J.X.) and the Scientific Research Foundation for High-level Faculty, China Pharmaceutical University (to J.X.). We thank Dr. Tongyu Liu for technical support with the bioinformatic analysis and Dr. Xin Chen (UCSF) for sharing the oncogene plasmids.

